# *Streptococcus pneumoniae* colonization modulates human nasal epithelial responses to respiratory syncytial virus infection

**DOI:** 10.64898/2026.02.05.703985

**Authors:** Leah A. Kafer, Isabel F. Escapa, Andrea I. Boyd, Alejandra Rivera Tostado, Amal Kambal, Sarah E. Blutt, Vasanthi Avadhanula, Pedro A. Piedra, Katherine P. Lemon

## Abstract

Respiratory syncytial virus (RSV) is a major cause of morbidity and mortality in infants globally. Specific nasal bacterial genera are differentially associated with RSV severity in infants: *Haemophilus* and *Streptococcus* with more severe disease and *Dolosigranulum* with healthy controls or milder outcomes. We hypothesized these differential bacterial effects begin at the epithelial level. Therefore, we established human nasal epithelial organoids differentiated at air-liquid interface (HNO-ALI) as a model system to assess effects of individual nasal microbionts on the epithelial response to subsequent RSV infection and of RSV on those microbionts. Infant-derived HNO-ALI were monocolonized with either *Streptococcus pneumoniae,* nontypeable *Haemophilus influenzae,* or *Dolosigranulum pigrum* one day before viral infection. RSV reduced colonizing *S. pneumoniae* and *D. pigrum* levels without affecting *H. influenzae. S. pneumoniae* precolonization uniquely reduced RSV levels during infection. *S. pneumoniae* precolonization also modulated the epithelial transcriptional response to RSV infection more so than *H. influenzae* or *D. pigrum*, with a pronounced effect on genes involved in immune response, cell cycle, stress, and growth signaling. Gene set enrichment analysis showed *S. pneumoniae* precolonization blunted RSV-induced increase in inflammatory and immune responses, consistent with *S. pneumoniae* also modulating RSV-induced cytokine production. Furthermore, *S. pneumoniae* precolonization blocked RSV-mediated dysregulation of cell-cycle genes, consistent with preventing arrest. Bacterial rescue of cell-cycle progression is a potential mechanism for reduced infectious virion production, since cell-cycle arrest enhances RSV replication. HNO-ALI facilitated elucidation of bacterial-viral-epithelial interplay at a frequent site of viral infection, directly linking nasal bacterial colonization to RSV infection dynamics.

**IMPORTANCE:** Most RSV-related hospitalizations occur in healthy young children, yet predictors of RSV severity in this population are limited. The nasal passages are a major microbiota site exposed to frequent respiratory viral infections. Specific nasal bacterial genera/species correlate with RSV clinical outcomes; however, the biological basis of these associations is poorly defined. Here, we established human nasal epithelial organoids differentiated at air-liquid interface (HNO-ALI) as a tractable experimental system for defining the interplay between bacteria, virus, and epithelial cells in the human nasal passages, enabling mechanistic insight into how the nasal microbiota contribute to RSV disease severity. Using this approach, we found that precolonization with each of the common nasal bacteria *S. pneumoniae*, *H. influenzae*, or *D. pigrum* differentially affected the nasal epithelial response to RSV infection. This work highlights that the nasal epithelium actively integrates microbial and viral signals and that variation in bacterial colonization can shape viral infection trajectories.

## INTRODUCTION

Respiratory Syncytial Virus (RSV) is a global threat to infant health, causing disease ranging from mild upper respiratory tract infection (URTI) to severe lower respiratory tract infection (LRTI), i.e., bronchiolitis and pneumonia (1). An estimated 33 million RSV-associated LRTIs occurred globally in children under 5 years old in 2019 with 3.6 million RSV-associated hospital admissions and up to 200,000 deaths (2). Infants hospitalized with RSV bronchiolitis are more likely to develop subsequent asthma and recurrent wheezing in the first decade of life (3). Although 98% of children are infected with RSV by the age of 2, only ∼10% develop LRTI (2, 4). Premature birth, congenital heart disease, and underlying pulmonary disease increase the risk of severe disease (5). However, previously healthy infants account for the majority of those hospitalized with RSV, with few predictors of disease severity in these children other than younger age (4, 6).

Variations in the composition of the bacterial nasal microbiota have emerged as a possible risk factor for severe RSV in healthy infants. Multiple studies show associations between the bacterial composition of the nasal microbiota and RSV infection and disease severity in infants (7–18). The nasal microbiota in most young children is enriched for one or two genera (17, 19–21). In cross-sectional, case-control, and prospective studies, profiles enriched for the bacterial genera *Haemophilus* and, to a lesser extent, *Streptococcus* are frequently associated with more severe RSV disease in infants (characterized by hypoxemia and the need for hospitalization), whereas nasal microbiota profiles enriched for the genera *Dolosigranulum* and/or *Corynebacterium* are associated with healthy controls or milder RSV outcomes (7–12, 15, 16, 18, 22–24). At the species level, the nasal pathobionts *Streptococcus pneumoniae* or *Haemophilus influenzae* are positively associated with increased RSV severity (9, 11, 12). In a prospective cohort study, the presence of residual respiratory symptoms two months after RSV infection was associated with a high relative abundance of *Haemophilus* and a lack of *Dolosigranulum* (16). These *in vivo* associations reflect correlations between nasal microbiota composition and RSV infection severity but do not establish causality or mechanism. Whether the viral infection alters bacterial colonization, the bacteria alter viral replication, the bacteria-virus dynamics modulate epithelial responses to infection, the immune cells play the dominant role, or all the above is still to be determined. Addressing these questions requires moving beyond correlational microbiome studies to experimental systems that enable mechanistic investigation of bacterial-viral-epithelial interplay.

A few animal studies have examined how bacterial precolonization affects RSV infection outcomes. In mice, nasal inoculation with *Dolosigranulum pigrum* 040417 prior to intranasal RSV infection reduces lung tissue injury, lung viral load, and modulates inflammation (25). In cotton rats, precolonization with some strains of *S. pneumoniae* prior to RSV infection increases viral load (26). Rodents provide a systemic model for RSV infection; however, unlike humans, they are only partially permissive for RSV (27). To address this limitation, the murine RSV analogue pneumonia virus of mice (PVM) has been used in murine models. When the nasal passages of infant mice are colonized with *S. pneumoniae* four days before PVM infection, the nasopharyngeal levels of *S. pneumoniae* increase and those of the virus PVM decrease (28). These informative animal models only partially recapitulate the symptomatology and pathophysiology of RSV infection in humans. Therefore, human-based experimental systems are also needed to mechanistically define how resident nasal bacteria influence RSV infection.

Human-based models have progressed from transformed/immortalized respiratory epithelial cell lines (which provided important mechanistic insights) to physiologically accurate well-differentiated primary human bronchial epithelial cells (pHBECs) (29, 30), to model RSV LRTI, and well-differentiated primary human nasal epithelial cells (pHNECs) (30–33), to model RSV URTI. Both of these differentiated primary epithelial cell models accurately recapitulate the response to RSV infection (31). However, only a small number of studies have used them to examine the bacterial-viral-epithelial interplay when bacteria colonization precedes RSV infection, which models the situation of the nasal microbiota, and these focus only on nasal pathobionts. For example, pre-exposing pHBECs to heat-inactivated nontypeable *H. influenzae* increases levels of proinflammatory cytokines during subsequent RSV infection (34, 35), whereas heat-inactivated *S. pneumoniae* does not, suggesting nontypeable *H. influenzae* might have a stronger effect on the tripartite dynamic (35). To our knowledge, no studies have examined live bacterial colonization effects on subsequent RSV infection using pHNECs, despite the nasal respiratory epithelium being the initial site of RSV infection and primary source for RSV transmission. Although differentiated pHBECs and pHNECs are excellent *ex vivo* models for RSV infection, each donation of primary nasal or bronchial epithelial cells has a finite lifespan of three-to-four passages under standard culture conditions, limiting use of the same genetic background for extensive mechanistic studies.

The recent development of nontransformed, nonimmortalized human airway epithelial organoids (36) followed by that of human nasal epithelial organoids differentiated at air-liquid interface (HNO-ALI) (37) overcomes the short-lifespan limitation of pHBECs/pHNECs. Each donated HNO line can be propagated, cryopreserved, and passaged repeatedly (prior to differentiation), serving as a long-term resource for repeated experimentation over the span of years (37–40). Derived from tissue-resident nasal stem cells and differentiated at an air-liquid interface, HNO-ALI are a remarkably accurate *ex vivo* model of the human nasal respiratory epithelium with a robust mucociliary blanket (37–40). Prior work from our groups show that HNO-ALI are an effective model to study either URT viral infection (37, 38, 40) or bacterial nasal colonization (39). Here, we combine these approaches to advance HNO-ALI as an *ex vivo* model system for investigating bacterial-viral-epithelial interplay. Because infant– and adult-derived HNO-ALI respond differently to RSV infection (38), and because clinical associations between specific nasal bacteria and RSV severity are predominantly observed in infants or young children, we exclusively used infant-derived HNO-ALI in this study.

Distinct nasal microbiota profiles are differentially associated with RSV infection severity *in vivo* in infants with *Haemophilus*– and *Streptococcus*-enriched profiles associated with more severe disease and *Dolosigranulum*-enriched profiles associated with healthy controls or milder outcomes (7–12, 15, 16, 18, 22–24). However, the mechanisms underpinning these associations are poorly defined. We hypothesized that these differential effects begin at the level of initial microbial-epithelial interplay on the nasal mucosa, where bacteria colonize and where RSV first infects. We used a reductionist approach modeling bacterial-viral-epithelial interplay with HNO-ALI to test the hypothesis that prior monocolonization with *S. pneumoniae*, nontypeable *H. influenzae*, or *D. pigrum* would have differential effects on RSV replication, bacterial levels, and the nasal epithelial response during RSV infection. Data from *in vivo* human microbiome suggests precolonization with nontypeable *H. influenzae* would have the strongest pro-inflammatory effect and *D. pigrum* the mildest. However, our results revealed unexpected dynamics. *S. pneumoniae* precolonization exerted the strongest effects with reduction in both bacterial counts and RSV infectious particles, while strongly modulating the epithelial transcriptional response to RSV. The latter included abrogating RSV-mediated dysregulation of cell-cycle-related transcripts, identifying a potential mechanism for the decrease in RSV infectious particles observed under *S. pneumoniae* precolonization, since cell cycle arrest is known to increase RSV titers (41, 42). This work demonstrates how infant-derived HNO-ALI can enable mechanistic dissection of bacterial-viral-epithelial relationships that underlie clinical associations between nasal microbiota composition and RSV disease severity.

## RESULTS

### RSV replicates in HNO-ALI at the nasal temperature of 34°C

Studies of RSV infection of pHBECs/pHNECs (26, 31, 43) and of HNO-ALI (37, 38, 40) are customarily conducted at the human internal body temperature of 37°C. However, the luminal temperature in the human nasal passages peaks at ∼34°C (44). Furthermore, nasal pathobionts, such as *Streptococcus pneumoniae* (45–47) and nontypeable *H. influenzae* (48), shift towards virulence at internal body temperature of 37°C compared to the lower temperatures of environmentally exposed epithelial surfaces (≤ 34°C). Based on these, we use 34°C to study bacterial nasal colonization of HNO-ALI (39). However, before using 34°C to study bacteria-RSV-nasal epithelial cell interplay, we first had to determine whether the globally circulating, contemporaneous RSV strain, RSV/A/Ontario (RSV/A/ON), infects infant-derived HNO-ALI at 34°C. We infected HNO-ALI with RSV at multiplicity of infection (MOI) of 0.01 (with a median of 6.6 x 10^3^ plaque forming units (PFUs) and 3.5 x 10^6^ gene copy number per HNO-ALI) (**Fig. 1A-B**). Apically, RSV levels peaked at 4 days post-viral infection (dpvi) with a median of 4.6 x 10^5^ PFUs, demonstrating productive replication of infectious viral particles at 34°C (**Fig. 1A**). As expected, neither RSV PFUs nor gene copies were detected in the basal medium, since RSV predominantly infects apical ciliated cells (38, 40, 49). Having shown that RSV replicates in infant-derived HNO-ALI at 34°C, we proceeded to investigate the tripartite interplay between nasal epithelial cells, RSV, and nasal bacteria implicated in variations in RSV infection severity.

**Figure 1.**
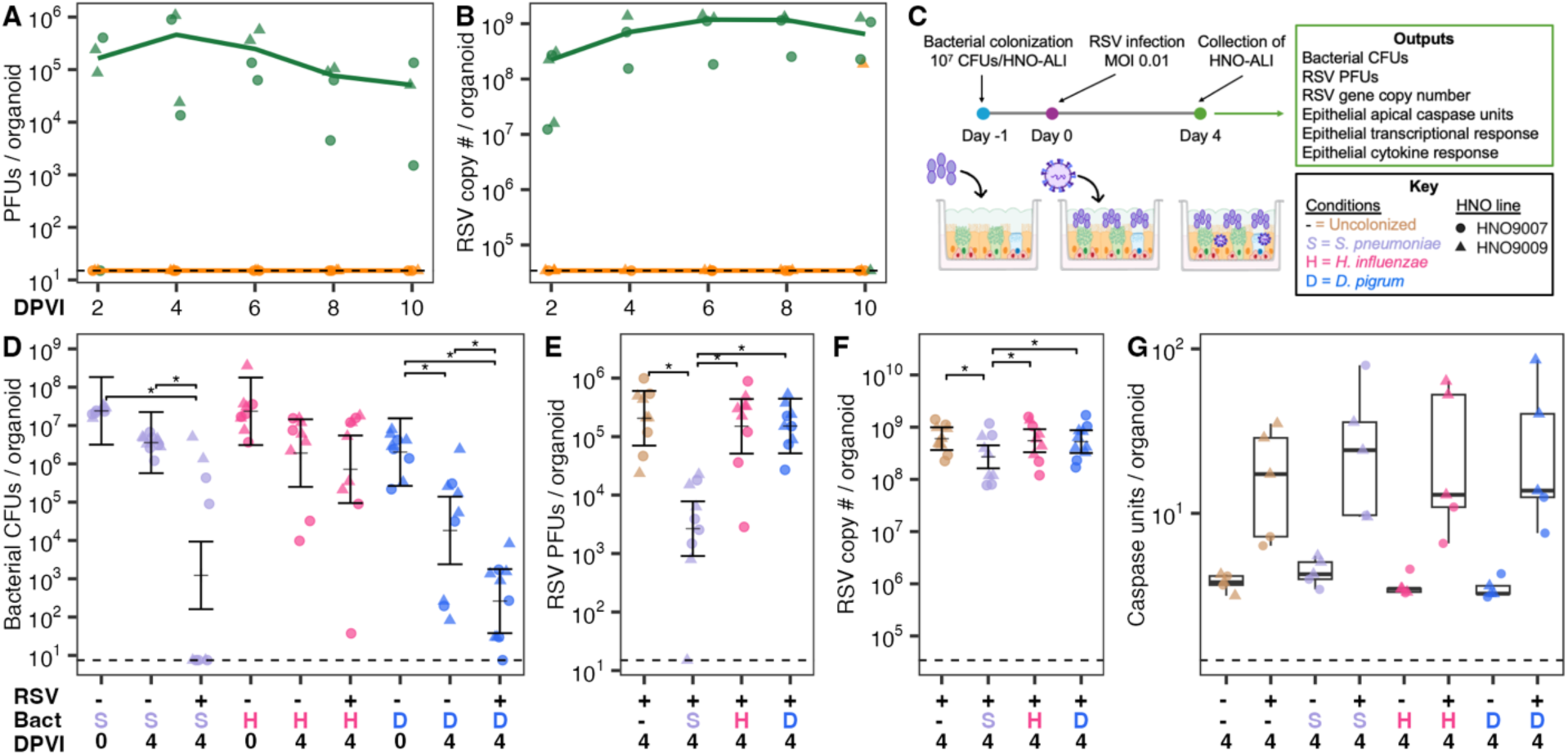
Precolonization of human nasal epithelium with *Streptococcus pneumoniae* reduces the level of RSV infectious particles. (**A**) PFUs and (**B**) gene copy number of virus recovered from HNO-ALI infected with RSV/A/Ontario at an MOI = 0.01 (median input of 6.6 x 10^3^ PFUs per HNO) show that RSV replicates at 34°C. Apical (green) and basal (orange) PFUs or gene copy numbers are shown over 10 days post-viral infection (dpvi). Lines connect the median at each timepoint. Two independent experiments in each of two HNO lines: HNO9007 (circles) and HNO9009 (triangles). (**C**) Experimental design for Figures 1D-G (as well as Figures 3 and 4) along with the color/symbol key. Infant-derived HNO-ALI were uncolonized (brown) or were monocolonized with *S. pneumoniae* (lavender S), nontypeable *H. influenzae* (pink H), or *D. pigrum* (blue D) at –1 dpvi followed by inoculation at 0 dpvi with RSV/A/Ontario at an MOI = 0.01 (median input of 6.2 x 10^3^ PFUs per HNO). Organoids were collected 4 dpvi to enumerate (**D**) bacterial CFUs, (**E**) RSV PFUs, (**F**) RSV gene copy number, and (**G**) caspase units per organoid. Horizontal brackets (D-F) indicate pairwise contrasts with adjusted *p* < 0.05. Vertical brackets (D-F) show the predicted mean values and 95% confidence intervals from the linear mixed-effects model (LMM). Data for D-G are from 5 independent experiments, 2 with HNO9007 (circles) and 3 with HNO9009 (triangles).

### RSV infection has differential effects on bacterial colonization of human nasal epithelial cells *ex vivo*

In infants, there are reports of the level of nasal *S. pneumoniae* or *H. influenzae* increasing during RSV infection, unlike for *D. pigrum* (16, 50). Epithelial cells, immune cells, inflammation, and RSV could all contribute to this. To determine whether nasal respiratory epithelial cells have a role in altering bacterial levels during RSV infection, we monocolonized HNO-ALI with either *S. pneumoniae,* nontypeable *H. influenzae,* or *D. pigrum* at ∼10^7^ colony forming units (CFUs) per transwell one day before RSV infection (i.e., –1 dpvi) (**Fig. 1C**). Bacteria were present on the HNO-ALI (**Fig. S1A**, **Table S1A**) on day 0 when HNO-ALI were either infected with RSV at 6.2 x 10^3^ PFUs per organoid (MOI = 0.01) or inoculated with buffer alone, then incubated at 34°C until 4 dpvi at which point bacterial CFUs were enumerated. Comparing RSV-infected to –uninfected HNO-ALI at 4 dpvi, *S. pneumoniae* CFUs decreased 2,913-fold with a bimodal distribution: there were no recoverable CFUs in 5 of 9 and a mean of 1.7 x 10^6^ CFUs in 4 of 9 of the RSV-infected HNO-ALI. In contrast, *H. influenzae* CFUs were similar with and without RSV infection. *D. pigrum* CFUs decreased 70-fold further with RSV versus without RSV infection (**Fig. 1D**, **Table S1A**). These data indicate that RSV/A infection of HNO-ALI can have differential species-level effects on members of the pre-existing bacterial nasal microbiota.

### Precolonization with *S. pneumoniae* reduces RSV levels during infection of human nasal epithelial cells *ex vivo*

Having shown that RSV infection has variable effects on the levels of colonizing nasal bacteria using HNO-ALI, we hypothesized that the bacterial species would have differential effects on RSV infection as well. To test the hypothesis that precolonization with bacteria affects RSV levels *ex vivo* in HNO-ALI, we compared RSV PFUs and gene copy number after infection of HNO-ALI alone compared with infection of HNO-ALI monocolonized in advance with *S. pneumoniae*, nontypeable *H. influenzae*, or *D. pigrum* (**Fig. 1C**). Only *S. pneumoniae* precolonization had a statistically significant effect on RSV PFUs and gene copy number, reducing PFUs 77-fold (**Fig. 1E**, **Table S1B**) and copy number 2-fold (**Fig. 1F**, **Table S1C**) compared to the uncolonized, RSV-infected controls. In contrast, *H. influenzae* and *D. pigrum* precolonization resulted in similar levels of RSV as in the uncolonized, RSV-infected HNO-ALI. RSV infection leads to apical cell apoptosis with leakage of intracellular enzymes from dying cells. As expected, RSV infection increased the level of caspase (an indicator of apoptosis) detected in apical washes. However, bacterial precolonization had no effect on apical caspase levels with or without RSV infection (**Fig. 1G**, **Table S1D**).

At 4 dpvi, only *S. pneumoniae* precolonization had decreased RSV PFUs (**Fig. 1E**, **Table S1B**). However, we hypothesized that this decrease started earlier in infection. To test this, we precolonized HNO-ALIs with *S. pneumoniae* for 24 h (1 d) before infecting with RSV, collecting independent HNO-ALI transwells at 1, 2, 3, and 4 dpvi to measure RSV PFUs and copy number (**Fig. 2**, **Table S1E-F**). One day of precolonization with *S. pneumoniae* decreased RSV PFUs as early as 1 dpvi, with a mean ∼4-fold decrease in RSV PFUs, followed by a 7-fold decrease at 2 dpvi, a 12-fold decrease at 3 dpvi, and a 31-fold decrease at 4 dpvi. Although *S. pneumoniae* precolonization decreased production of infectious RSV particles (**Fig. 2A**, **Table S1E**), there was no significant decrease in RSV gene copy number from 1-to-4 dpvi (**Fig. 2B**, **Table S1F**). However, PCR is very a sensitive method that can detect both defective viral particles and degrading viral genomes.

**Figure 2.**
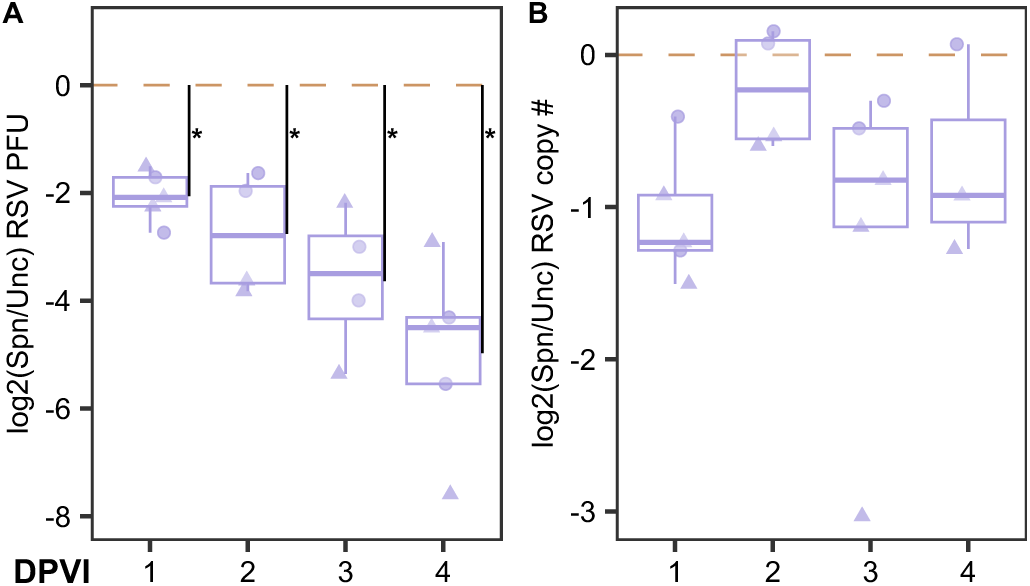
*S. pneumoniae* precolonization decreases levels of RSV infectious particles over time compared to uncolonized HNO-ALIs. Log_2_ fold-change of RSV levels in HNO-ALI colonized with *S. pneumoniae* compared to uncolonized HNO-ALIs across four dpvi. (**A**) RSV PFUs and (**B**) gene copy numbers. Data are from 5 independent experiments, 2 with HNO9007 (circles) and 3 with HNO9009 (triangles). Boxplots show median and interquartile range. Vertical lines and asterisks indicate time points at which *S. pneumoniae* significantly decreased RSV infectious particles relative to the uncolonized control with adjusted *p* < 0.05. Dashed brown line indicates baseline if RSV levels in colonized and uncolonized HNO-ALI were the same.

### *S. pneumoniae* colonization modulates the HNO-ALI transcriptional response to RSV infection more so than does nontypeable *H. influenzae* or *D. pigrum*

Human *in vivo* microbiome studies associate these bacteria with differing levels of RSV infection severity (7–12, 15, 16, 18, 23). Based on this, we hypothesized that precolonization with each bacterium would differentially affect the epithelial response to RSV infection. To test this, we examined gene expression profiles in HNO-ALI precolonized for one day prior to RSV infection with either *S. pneumoniae*, *H. influenzae*, or *D. pigrum*, alongside an uncolonized control. Principal component (PC) analysis (**Fig. S2A**) showed that RSV infection was the dominant driver of variation in transcript levels, corresponding to the separation observed along PC1 (81% of variance), while PC2 (9% of variance) primarily reflected HNO donor line. This strong antiviral signature is consistent with previous reports of robust epithelial responses to RSV (31, 40, 51–54). The effects of bacterial colonization were less pronounced, indicating that colonization does not override the host response to viral infection but might have a modulatory effect. To capture this, we focused our statistical analysis on the bacteria-virus interaction terms, which assess whether the HNO-ALI transcriptional response to RSV infection differs depending on bacterial colonization status (rather than simply comparing RSV plus each bacterium to RSV alone). Differential expression analysis revealed markedly different transcriptional modulation of the response to virus across bacterial species (**Fig. 3A**, **Tables S2-4**). *S. pneumoniae* precolonization had the most pronounced interaction effect on the epithelial response to RSV infection, with 597 statistically significant increased and 1028 decreased transcripts, compared to 6 increased and 23 decreased transcripts with *H. influenzae* and 36 increased and 161 decreased transcripts with *D. pigrum*. This analysis revealed that *S. pneumoniae* exerts the strongest influence on reshaping the nasal epithelial cell transcriptional response during RSV infection.

**Figure 3.**
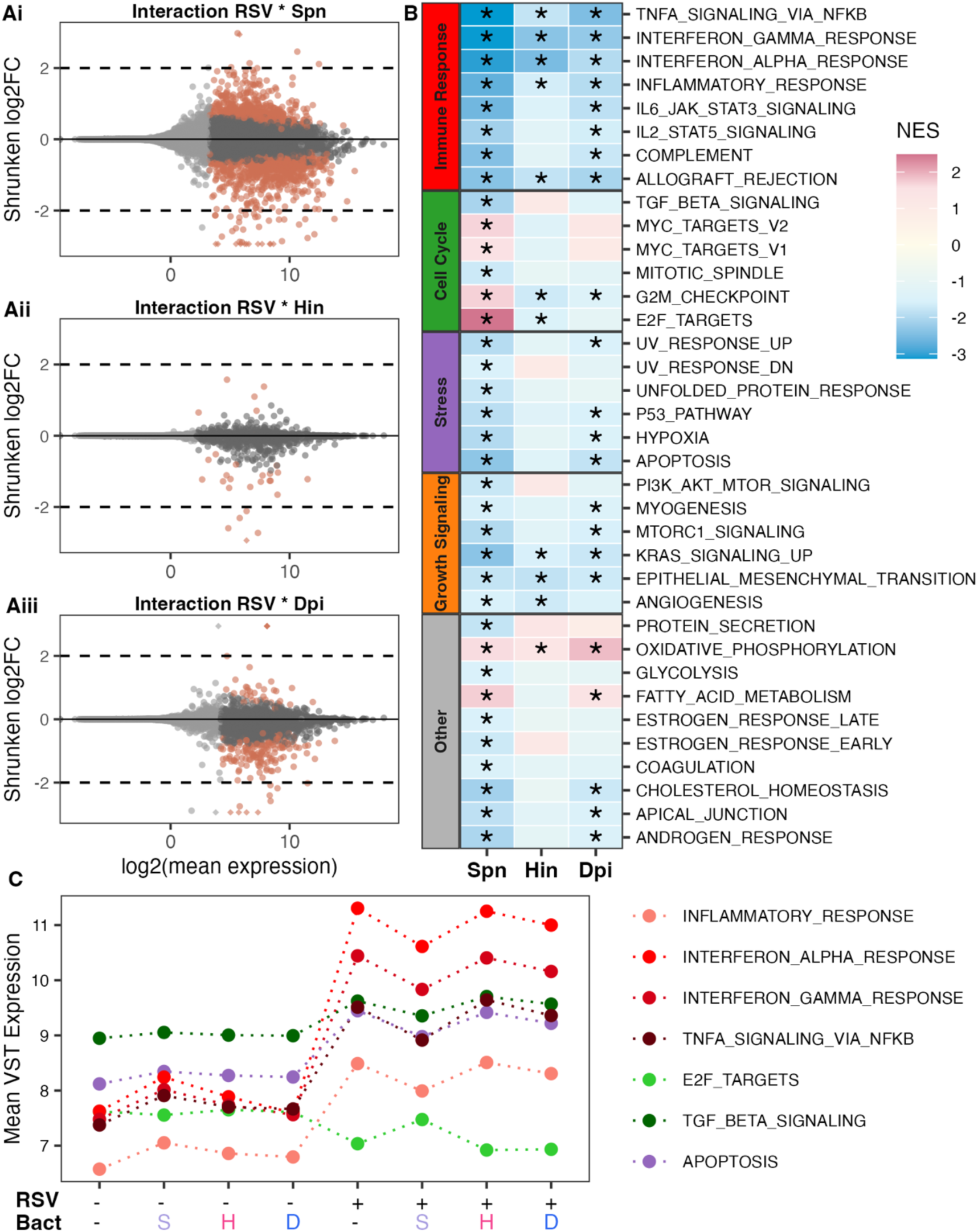
*S. pneumoniae* precolonization modulate the HNO-ALI response to RSV compared to the uncolonized HNO-ALI response to RSV more so than nontypeable *H. influenzae* and *D. pigrum*. (**A**) MA plots showing differential gene expression for bacteria-virus species interaction terms. Points represent individual genes plotted by log2 mean expression (x-axis) and shrunken log2 fold change (y-axis). Brown points indicate significantly differentially expressed genes (FDR *p* ≤ 0.05). Gray points represent genes that did not meet those thresholds, with light gray representing genes where *p*-values were not calculated. Horizontal dashed lines mark the ±2 log2FC cutoff. Positive log2FC values indicate genes where the virus effect is stronger in the given species compared to uncolonized conditions, while negative values indicate a weaker effect. Y-axis is constrained to ±3 for visualization, with outliers clipped to the boundary and represented as diamonds. (**B**) Gene Set Enrichment Analysis (GSEA) heatmap showing Normalized Enrichment Score (NES) for MSigDB Hallmark gene sets across the same bacteria-virus interaction contrasts. Gene sets are grouped by biological category and filtered to include only those with FDR *p* < 0.05 and |NES| ≥ 1.5 in at least one contrast. Color gradient represents NES values and asterisks indicate statistical significance. Gene sets are ordered by their dynamic range across contrasts, with the most variable responses at the top. (**C**) Mean Variance Stabilizing Transformation (VST)-normalized expression trajectories for selected gene sets from panel B. For each gene set, leading-edge genes (the core subset of genes that contributed most to the enrichment signal) were obtained from GSEA results and their VST-normalized expression values averaged per condition. Each colored line represents a distinct gene set, showing coordinated expression changes across bacterial colonization and RSV infection conditions. The – = uncolonized; S/Spn = *S. pneumoniae* (purple); H/Hin = nontypeable *H. influenzae* (pink); D/Dpi = *D. pigrum* (blue).

To identify the biological pathways underlying these differential effects of bacterial colonization on the response to RSV, we performed Gene Set Enrichment Analysis (GSEA) using Hallmark gene sets on bacteria-virus interaction contrasts (i.e., genes where the RSV response differs by bacterial colonization status). GSEA reports a Normalized Enrichment Score (NES), a size-adjusted measure of coordinated up– or down-regulation within a gene set. Our analysis revealed coordinated changes across multiple functional categories in how bacterial precolonization modulated RSV-induced transcriptional responses. Notably, *S. pneumoniae* precolonization most substantially changed how HNO-ALI responded to RSV, with significant effects (FDR p < 0.05 and |NES| ≥ 1.5) across immune response, cell cycle, stress, and growth signaling gene sets compared to the other two bacteria (**Fig. 3B**, **Table S5A**).

*S. pneumoniae* precolonization attenuated (strong negative NES) the transcriptional response to RSV infection of several immune response gene sets (**Fig. 3B**, red category). This pattern was particularly evident across the following immune response gene sets: “inflammatory response”, “interferon-alpha response”; “interferon-gamma response”, and “TNF-alpha signaling via NFκB”. This was observed in a much more subtle manner for *H. influenzae* and *D. pigrum*. Mean expression trajectories for leading-edge genes driving this enrichment revealed that *S. pneumoniae* colonization alone (without RSV infection) elevated baseline expression of inflammatory and interferon response genes relative to uncolonized controls (**Fig. 3C**, red trajectories, **Table S5B**, **Fig. S2Bi-iv**). However, upon RSV infection, these same pathways showed substantially blunted induction in *S. pneumoniae*-colonized HNO-ALI compared to the robust induction in uncolonized HNO-ALI.

Cell cycle-related pathways also showed a striking modulatory pattern due to *S. pneumoniae* precolonization of HNO-ALIs (**Fig. 3B**, green category). The mean expression trajectory for “E2F targets” leading-edge genes showed that after RSV infection, the cell cycle response diverged based on the colonizing bacterium (**Fig. 3C**, light green, **Table S5B**, **Fig. S2Bv**). RSV infection of uncolonized HNO-ALI, as well as *D. pigrum*– and *H. influenzae*-precolonized HNO-ALI, caused a pronounced decrease in transcript levels of E2F targets. This is consistent with previous reports of RSV-induced cell cycle arrest mediated by TGF-β signaling (42). In contrast, RSV-infected *S. pneumoniae*-precolonized HNO-ALI had E2F target transcript levels equivalent to those of uninfected, uncolonized HNO-ALI, indicating that *S. pneumoniae* colonization prevented RSV suppression of these cell-cycle-associated genes. Notably, transcripts related to “TGF-β signaling” (**Fig. 3C**, dark green, **Table S5B**, and **Fig. S2Bvi**) were also suppressed in *S. pneumoniae*-precolonized HNO-ALI after RSV infection compared to the other bacteria and uncolonized controls.

### *S. pneumoniae* precolonization modulates HNO-ALI cytokine response to RSV infection

Transcriptomic analysis revealed that bacterial precolonization, particularly with *S. pneumoniae*, modulated RSV-induced expression of many genes involved in inflammation/immunity (**Fig. 3**). Based on this, we tested whether bacterial precolonization affected infant-derived HNO-ALI cytokine production in response to RSV infection at 4 dpvi with a multiplex panel of 39 cytokines. We collected apical measurements to model cytokines released into nasal mucus, and therefore present at the site of bacterial colonization and viral infection, as well as basal measurements to model cytokine release into tissue and the circulation, from whence immune cells are recruited in the host. PC analysis accounting for batch effects indicated RSV infection status again corresponded to the largest source of variation with samples separating clearly along PC1 (explaining >45% of the variance), regardless of bacterial colonization status (**Fig. S3**). Partial least squares discriminant analysis (PLS-DA, **Fig. 4A-B**) confirmed clear separation of RSV-infected vs. –uninfected samples in component 1, (∼46% apical, ∼64% basal variance) with additional separation by *S. pneumoniae* precolonization (purples) along component 2 (∼10% variance in both compartments).

**Figure 4.**
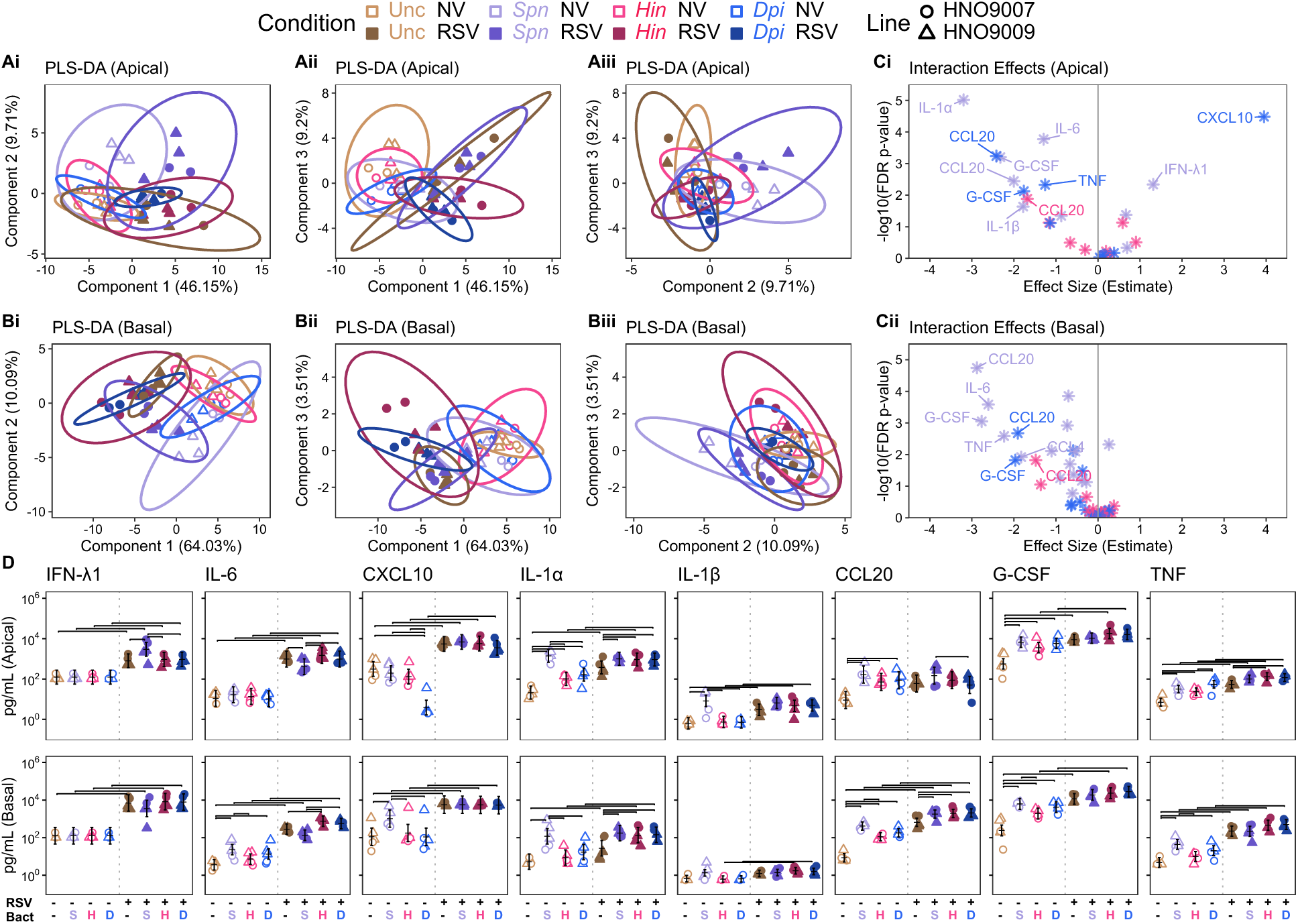
*S. pneumoniae* precolonization differentially modulates the human nasal epithelial cytokine response to RSV infection at four dpvi compared to *H. influenzae* and *D. pigrum*. PLS-DA score plots show that RSV infection (filled shapes) had a large impact on HNO-ALI cytokine profiles in the (**A**) apical wash or (**B**) basal medium. Modulation of this response by *S. pneumoniae* precolonization is visible in component 2 (i and iii), in contrast to *D. pigrum* or *H. influenzae*. PLS-DA was performed using the optimal number of components selected via cross-validation (**Fig. S3**). Points represent individual samples colored by bacterial treatment in RSV infected (filled, dark colors) or uninfected (open, light colors) HNO-ALIs, with shape representing HNO-ALI line. Ellipses represent 95% confidence intervals around group centroids. Analysis was performed accounting for experimental date as a repeated measure (multilevel design) with data mean-centered and scaled to unit variance prior to analysis. (**C**) Volcano plots of bacteria-virus interaction effects estimated by an LMM comparing the virus effect in each bacterial treatment versus in the uncolonized (no bacteria) control for (i) apical washes and (ii) basal medium. Points represent cytokines with the estimated interaction effect size on the x-axis and the statistical significance (-log10 FDR *p*) on the y-axis. Labeled cytokines met the threshold for both significance (FDR *p* < 0.05) and effect size (|estimate| ≥ 1.25). (**D**) Levels (pg/ml) of the HNO-ALI produced cytokines (pg/mL) from C that were detected in the apical wash (upper row) or the basal medium (lower row). Data are from five independent experiments, two with HNO9007 (circles) and 3 with HNO9009 (triangles). Samples colored by bacterial treatment in RSV infected (filled, dark colors) or uninfected (open, light colors) HNO-ALI, with shape representing HNO line and – = uncolonized; S = *S. pneumoniae* (purples); H = nontypeable *H. influenzae* (pinks); D = *D. pigrum* (blues).

Statistical analysis of bacteria-virus interaction effects identified specific cytokines whose response to RSV infection differed by bacterial precolonization (**Fig. 4C**, **Table S6A-B**). The levels (pg/ml) of some of these cytokines showed statistically significant variability across bacterial conditions in either or both RSV-infected and –uninfected HNO-ALI, as discussed in detail next (**Fig. 4D**, **Table S6A-B**).

IFN-λ1 is a type 3 interferon (mucosal interferons) and higher nasopharyngeal levels are associated with decreased likelihood of hospitalization in RSV-infected infants (55). During RSV infection on HNO-ALI, *S. pneumoniae* precolonization increased apical IFN-λ1 by ∼4-fold compared to uncolonized HNO-ALI, whereas *D. pigrum* and *H. influenzae* had no effect. None of the bacteria influenced IFN-λ1 release into the basal medium.

IL-6 has pleiotropic effects (stimulating immune cell proliferation and differentiation to resolve infections) with an important role in acute phase response (56–58). However, high nasopharyngeal levels can cause excessive inflammation and are associated with increased RSV severity (59, 60). Apically, *S. pneumoniae* precolonization reduced IL-6 during RSV infection 2-to-3-fold compared to *H. influenzae*, *D. pigrum,* or no precolonization (**Fig. 4D**, **Table S6A-B**). Basally, *S. pneumoniae* colonization increased IL-6 by ∼8-fold compared to uncolonized HNO-ALI but reached levels similar to uncolonized HNO-ALI during subsequent RSV infection. *D. pigrum* increased IL-6 4-fold without RSV but not with RSV and *H. influenzae* had no effect without RSV but increased it 3.3-fold with RSV.

CXCL10 is a pleiotropic cytokine implicated in both antiviral and antibacterial responses that acts by binding to the CXCR3 receptor, which then has a multitude of downstream effectors (61). Higher nasopharyngeal levels of CXCL10 are associated with decreased hospitalization risk in RSV-infected infants (55, 62). The bacteria alone had differential effects by 4 dpvi. In fact, *D. pigrum* precolonization reduced apical production of CXCL10 prior to RSV infection (**Fig. S1B-C**, **Table S6C-D**), which is similar to what we observed previously in adult-derived HNOs (39). However, RSV infection elevated CXCL10 similarly across all conditions in both compartments, overriding the effects of the bacteria.

Higher levels of inflammasome-related IL-1β in nasopharyngeal samples from children are associated with increased severity in RSV infection (59, 60). Damaged/stressed mammalian cells release the alarmin IL-1α, which then binds to and activates the IL-1 receptor leading to release of IL-18 and IL-1β (63). *S. pneumoniae* precolonization of uninfected HNO-ALI increased both IL-1α and IL-1β (**Fig. 4D**, **Table S6A-B**). *S. pneumoniae* increased apical IL-1α 66-fold, compared to ∼ 5-fold with *D. pigrum* or *H. influenzae,* and alone increased basal IL-1α (∼24-fold). *S. pneumoniae* alone also increased apical IL-1β (∼13-fold). However, subsequent RSV infection increased the apical and basal levels of both cytokines in all three bacterial conditions to approximately that of the *S. pneumoniae* precolonized alone, indicating that the epithelial response to RSV infection dominated.

CCL20 is a chemokine that attracts lymphocytes (64) and dendritic cells (65). G-CSF is a growth factor that enhances neutrophils survival and modulates their epithelial activity (66). TNF is a pleiotropic pro-inflammatory cytokine that recruits innate immune cells to fight off infection (67), and whose overproduction can lead to prolonged inflammation and increased tissue damage in RSV infection of young children (59). In uninfected HNO-ALI, all three bacterial species increased CCL20, GCSF, and TNF after 5 days (i.e., 4 dpvi), with *S. pneumoniae* generally inducing the largest effects, followed by *D. pigrum*, then *H. influenzae* (**Fig. 4D**, **Table S6A-B**). In contrast, RSV infection induced an increase in all three of these cytokines apically and basally compared to the uninfected HNO-ALI regardless of colonization status, with the exception of apical CCL20.

These data demonstrate that RSV has the predominant effect on cytokine production by HNO-ALI during viral infection and that bacterial precolonization can modulate this response, with *S. pneumoniae* doing so the most (**Fig. 4C**).

## DISCUSSION

Overall, this study demonstrates that human nasal epithelial cells actively integrate microbial and viral signals and that variation in pre-existing bacterial colonization can shape viral infection trajectories. We used infant-derived HNO-ALI to mechanistically investigate epidemiological reports that distinct nasal microbiota profiles are differentially associated with RSV infection severity in infants: *Haemophilus* and *Streptococcus* with more severe infection and *Dolosigranulum* with healthy controls or milder infection (7–12, 15, 16, 18, 22–24). Precolonization of HNO-ALI with each of the major nasal species from these genera had differential effects on RSV infection outcomes. *S. pneumoniae* had the strongest effects with statistically significant reduction in bacterial and viral counts (**Fig. 1D-E**) as well as modulation of epithelial transcription (**Fig. 3**) and cytokine production (**Fig. 4**). In contrast, *H. influenzae* had the mildest effects with stable bacterial and viral counts and more modest modulation of epithelial transcription and cytokine production. The effect of *D. pigrum* precolonization on nasal epithelial cell responses to RSV infection was generally in between that of the two pathobionts with reduction of bacterial counts, no effect on viral counts, and moderate modulation of epithelial transcription and cytokine production.

Precolonization of HNO-ALI with *S. pneumoniae* caused a decrease in infectious RSV viral particles (**Fig. 1E**) that was evident as early as one dpvi (**Fig. 2**). Similar findings are reported in mice, where nasal colonization with *S. pneumoniae* before nasal infection with the murine RSV analogue PVM decreases virus levels (28), including with strain 603 (serotype 6B), which we also used here. This reduction in viral particles appears paradoxical given that, in young children, nasal microbiota enriched for *S. pneumoniae* are more often positively associated with increased RSV infection severity (9, 15, 16, 22, 23). However, higher levels of detectable nasal RSV in infants are associated with decreased disease severity (68), suggesting the possibility that *S. pneumoniae* reduction in RSV PFUs in nasal epithelial cells might contribute to increased disease severity.

A potential mechanism for the *S. pneumoniae*-mediated reduction in RSV particles is suggested by our finding that only *S. pneumoniae* precolonization prevented the RSV-mediated dysregulation of cell-cycle gene transcripts (**Fig. 3B-C**). This RSV-mediated dysregulation pattern is consistent with cell cycle arrest, a known consequence of RSV infection in pHBECs. However, reports conflict on whether arrest occurs in G0/G1 or in G2/M, as well as the underlying mechanisms (42, 69). Viral-induced cell cycle arrest correlates with an increase in viral reproduction (41, 42), suggesting that *S. pneumoniae*’s abrogation of RSV-mediated dysregulation of cell-cycle-associated transcripts (**Fig. 4C**) might directly contribute to the observed decrease in RSV infectious particles (**Fig. 1E**). Very little is known about the effect of *S. pneumoniae* on epithelial cell cycle. Pneumolysin alone is reported to induce G2/M cell cycle arrest (70); however, the findings here point to live *S. pneumoniae* cells preventing cell cycle arrest. In pHBECs, RSV infection increases TGF-β secretion, leading to cell cycle arrest (42). Based on our findings, it is possible that *S. pneumoniae* precolonization might disrupt the RSV-induced cell cycle arrest program in respiratory epithelial cells through suppression of TGF-β signaling (**Fig. 3C**). This is an avenue for future research after first determining the specific cell cycle stage at which arrest occurs in HNO-ALI.

The mildest effects were observed with nontypeable *H. influenzae* precolonization, which was unexpected given that nasal microbiota enriched for *Haemophilus* are often positively associated with increased RSV infection severity (8–12, 15, 16, 18, 23, 24). However, we found no evidence for mutual influence between nontypeable *H. influenzae* and RSV at the level of epithelial cells using infant-derived HNO-ALI, with neither’s levels affected by the other (**Figs. 1D-E**). Similarly, precolonizing HNO-ALI with nontypeable *H. influenzae* had only a modest effect on the epithelial transcriptional response to RSV, with the lowest number of differentially expressed genes (**Fig. 3Aii**) and of statistically significant gene sets in the bacteria-virus statistical interaction analysis (**Fig. 3B**). It also showed the fewest cytokine-level effects attributable to the bacteria-virus interaction term among the three bacterial species (**Fig. 4C**). These results leave the impression that, although the nasal epithelial cells detect nontypeable *H. influenzae* colonization, it is more evasive than the other two bacteria. These results raise the possibility that tissue-resident immune cells, rather than epithelial responses per se, might be a key component in the relationship between *Haemophilus* and RSV infection severity. A possibility that warrants study in a more complex model inclusive of immune cells.

Nasal microbiota enriched for *Dolosigranulum* (most probably *D. pigrum*) are frequently associated with healthy controls or decreased severity of RSV and other respiratory viral infections in infants (7, 16, 18, 71–73). In mice, nasal inoculation with *D. pigrum* 040417 before intranasal RSV infection reduces viral load in the lungs, lung injury, and modulates inflammation; however, nasal effects are not reported (25). To our knowledge, this is the first study examining *D. pigrum* colonization effects on RSV infection in an *ex vivo* model of infant nasal respiratory epithelium. Similar to adult-derived HNO-ALI (39), *D. pigrum* monocolonized infant-derived HNO-ALI at lower levels than either pathobiont, and RSV infection further decreased its levels (**Fig. 1D**), suggesting this potential mutualist is more susceptible to epithelial defenses. Although it had no effect on RSV PFUs or copy number (**Fig. 1E-F**), *D. pigrum* did have some modulatory effect on the transcriptional response of HNO-ALI to RSV infection (**Figs. 3Aiii**). For example, similar to *S. pneumoniae*, *D. pigrum* attenuated (negative NES) the transcriptional response of several immune response gene sets to RSV infection, though to a lesser degree (**Figs. 3B**). Although this dampening of the inflammatory response by *D. pigrum* aligns with its clinical association with less severe RSV disease, understanding how this bacterial-viral-epithelial interplay translates into clinical outcomes requires further investigation.

Study limitations include use of a single strain of each of the three bacterial species and of the virus, use of two HNO donor lines, and a reductionist tripartite model without immune cells. Prior studies suggest serotype– or strain-specific variability in the relationship between *S. pneumoniae* and RSV (74). Although RSV/A and RSV/B strains cause a similar disease phenotype in infants, it is unknown whether *S. pneumoniae* would similarly modulate RSV/B infection of HNO-ALI. In using two different infant-derived HNO-ALI lines, we opted for repeated independent experiments with the same lines as part of rigorously developing this tripartite model system. Going forward, this experimental model could be applied to additional HNO-ALI lines to assess for effects of host genetic variation; to additional bacterial and viral strains and species to assess for strain– and phylogenetic-based microbial variations; and to the addition of immune cells to the epithelial layer to assess the interplay between bacteria, virus, epithelial cells, and immune cells.

The HNO-ALI model is well-suited for future testing of specific genetic mutants in both the microbes and the HNO donor lines to further define molecular mechanism. This approach is already being used in human intestinal organoids (75). More broadly, this system is adaptable to other upper respiratory tract viruses and bacteria using infant– or adult-derived HNO-ALI, enabling systematic investigation of how microbiota composition shapes viral infection outcomes across the lifespan. HNO-ALI are a tractable experimental framework for translating correlative microbiome findings into causal, mechanistic understanding of how the microbiota modulates viral infections.

## MATERIALS and METHODS

### Microbial strains and HNO lines

RSV/A/USA/BCM813013/2013 (genotype Ontario) (RSV/A/ON) (76) was used for all infections at an MOI of 0.01, which was calculated based on the number of epithelial cells per HNO for each line and experimental date. The following bacterial strains were used: *Dolosigranulum pigrum* strain KPL3065 (77) and *Streptococcus pneumoniae* strain 603 (aka GA03212) (78), and nontypeable *Haemophilus influenzae* strain 86-028NP (79). Two infant-derived HNO lines were used: HNO9007 and HNO9009 (38).

### HNO-ALI

Isolation, propagation, and differentiation of HNO-ALI was done similarly to our previous description (39), with the following important changes in the differentiation protocol that have enhanced the reliability of the organoid cultures: HNO-ALI monolayers were seeded at a density of 85,000 – 100,000 cells per Transwell® insert (Corning #3470), and apical airway organoid (AO) medium was removed 2-to-3 days after seeding. (Previously, HNO-ALI monolayers were seeded at a density of 250,000 cells per transwell, and AO medium was removed 4 days after seeding (39).)

### Additional HNO-ALI Quality Controls

As an additional layer of quality control, we measured the transepithelial electrical resistance (TEER) to assess barrier integrity one-day prior to experimentation. The minimum TEER measurement of the HNO-ALI for each experiment was 600 Ohms. The TEER was measured for one representative transwell of HNO-ALI per line ∼24 h before the start of each experiment (date), since the required addition of apical buffer for measurements disrupted the air interface. As we previously described (39), we added 100 µL of Earle’s Balanced Salt Solution (EBSS) without calcium, magnesium, or phenol red (Gibco 14155063) to the apical side of the selected HNO-ALI, placed the TEER meter probe (World Precision Instruments EVOM with STX4 electrodes) into the liquid of the transwell per (80), and recorded the ohms. The median TEER for each line across all experiments was as follows: HNO9007 1140 ohms, HNO9009 1175 ohms. (The range of the average TEER (for measured wells) per experiment was as follows: HNO9007 915-1390 Ohms, HNO9009 830-1370 Ohms.) After confirming a TEER ≥ 600 ohms from one HNO, we changed the basal medium in all HNO-ALI wells to AO differentiation medium (AODM) without heparin or hydrocortisone (AODM w/o HH) and placed the 24-transwell plate into a humidified 5% CO_2_ incubator at 34°C.

Also, for added quality control, we confirmed that each representative HNO-ALI used for TEER measurement was negative for *Mycoplasma,* as we previously described (39). After removing the TEER probe, HNO-ALI cells were scraped from the transwell with a 200 µL pipet tip, resuspended in 100 µL of EBSS, and transferred to a clean 1.5 mL microcentrifuge tube for mycoplasma testing. Basal medium from that well was also collected into a separate 1.5 mL microfuge tube for mycoplasma testing. Each microfuge tube was then heated at 95°C for 15 minutes (min), allowed to cool for 5-10 min, then vortexed, before removing 2 µL for PCR reactions. Each sample underwent a separate PCR reaction with the Biovision Mycoplasma PCR Detection kit, per manufacturer’s protocol. To detect samples positive for *Mycoplasma*, we performed electrophoresis on the PCR reactions using a 1% TopVision agarose Tris Acetate EDTA (TAE) gel and then visualized any DNA with SYBR Safe (Invitrogen #S33102).

### Enumeration of epithelial cells per organoid transwell

We enumerated cells per HNO-ALI (as previously described (39)) in order to calculate the RSV MOI. In brief, 200 μL of 0.25% Trypsin-EDTA (Gibco #25200056) was added to an HNO-ALI transwell and incubated at 37°C in a humidified 5% CO_2_ incubator for 10-15 min. The HNO-ALI was then gently pipetted up and down 3 times with a 200 μL pipet tip, followed by incubation for another 10-15 min. To separate the nasal epithelial cells from each other, they were gently pipetted up and down ≥ 20 times before transfer to a 1.5 mL microfuge tube. A cell aliquot was mixed with equal parts trypan blue to count viable cells with a BioRad TC20 automated cell counter.

### RSV infection time course

On day 21 of differentiation, HNO-ALI basal medium was replaced with AODM w/o HH and HNO-ALI were placed at 34°C in a humidified 5% CO_2_ incubator for 24 h. After 24 h (on –1 dpvi), 15 µL of EBSS was added to the apical side of each HNO-ALI to parallel the protocol for the negative control during bacterial precolonization (see below). The 24-well transwell plates were centrifuged in an Eppendorf 5430R at 120 x g for 1 min at RT prior to incubation for 23 h more. On the day of viral infection (day 0), the basal medium was changed by transferring the transwells into a new plate with 600 µL of fresh AODM w/o HH per well. The spent medium was collected and frozen at –80°C for downstream assays. HNO-ALI were incubated for 1 h at 34°C in a humidified 5% CO_2_ incubator before infection with RSV/A/ON (MOI 0.01). The viral stock was diluted in EBSS to achieve the correct MOI and 15 µL was pipetted onto the apical side of each HNO-ALI, preserving the ALI. The uninfected control HNO-ALI received 15 µL of EBSS alone. HNO-ALI were incubated at 34°C in a humidified 5% CO_2_ incubator for 2, 4, 6, 8, or 10 days. One infected HNO-ALI was collected at each timepoint; the spent basal medium from each harvested HNO-ALI was collected into a 1.5 mL microcentrifuge tube and frozen at –80°C to be used for cytotoxicity assays. Remaining HNO-ALI received fresh basal medium every 48 h until their designated collection timepoint by transferring the transwells to a new plate with fresh AODM w/o HH. To collect virus, 100 µL of Gibco 0.25% trypsin-EDTA, phenol red (ThermoFisher 25200) was added to the apical side of an HNO-ALI and incubated for 15 min at 34°C in a humidified 5% CO_2_ incubator before addition of 50 µL of AODM. The bottom of the transwell was scraped with a 200 µL pipet tip prior to pipetting up and down 30-40 times to remove the cell layer. The resuspended cells were transferred to a microcentrifuge tube with 150 µL of AODM w/o HH and mixed 30 times to create a homogenous suspension. We then transferred 150 µL of this suspension to a second microfuge tube containing 150 µL of Iscove’s Modified Dulbecco’s Medium (IMDM, Gibco 12440053) with 15% glycerol for plaque assays. The remaining 150 µL in the first tube was used for RNA extraction and reverse-transcription quantitative PCR (RT-qPCR). All tubes were flash frozen in an ethanol-dry ice bath and stored at –80°C until use.

### Viral copy number measurement

Frozen HNO-ALI wash samples designated for RNA extraction and RT-qPCR were thawed at RT and extracted using the QiaAmp Viral RNA mini kit per the manufacturer’s instructions. RT-qPCR was performed using the Ag-PATH ID kit as previously described (68) and a Biorad CFX Opus 384. Serial dilutions of ultramers of the RSV/A N gene were assayed with each set of samples to generate a standard curve and determine the RSV copy number in each sample. The code for this calculation can be found at https://klemonlab.github.io/RSVBac_Manuscript.

### Infectious virion measurement

Similar to previously described (37, 38), a frozen aliquot of HEp-2 cells was removed from liquid nitrogen, thawed on ice, and resuspended in antibiotic-free HEp-2 medium, consisting of 10 mL of MEM (Corning 10010CM) with 10% defined fetal bovine serum (DFBS) (Cytiva SH30070.01) and 1% L-glutamine (Gibco 25030081). Cells were then centrifuged at 120 x g for 10 min at RT (Eppendorf 5702R), and the supernatant was removed. Cells were resuspended in 10 mL of antibiotic-free HEp-2 medium and split into two 25-cm^2^ flasks, adding medium to bring the volume of each flask to 7-10 mL total. Flasks were incubated at 37°C with 5% CO_2_ for approximately 2 days or until the cells were 80-90% confluent. Cells from one flask were split into two 75-cm^2^ flasks and allowed to grow until 80-90% confluent. With the other flask, the cells were scraped off the bottom of the flask with a serological pipet and mixed into the medium, then 1 mL was transferred to a microfuge tube to be used in a mycoplasma assay. Only cells with a negative mycoplasma test were used for plaque assays. The 75-cm^2^ flasks were then used to seed 24-well plates at 1-2 x 10^5^ cells/mL in each well and allowed to grow until 80-90% confluent in a 37°C incubator with 5% CO_2_, prior to use for plaque assays. The viral plaque assays were completed as detailed in (81). Briefly, the samples designated for plaque assays were thawed at RT and each was serial diluted from 10^-1^ to 10^-4^ in HEp-2 infection medium (MEM with 2% DFBS,1% Antibiotic-Antimycotic (AA, Gibco 15240062), and 1% L-glutamine). In duplicate, 200 µL of each dilution was added to one well of the 24-well plate by first removing the medium on the HEp-2 cells then adding the sample dilution. Plates were swirled to distribute the viral suspension, then incubated at 37°C with 5% CO_2_ for 1.5-to-2 h. The viral suspension was removed from each well and 1.5 mL of plaque medium (1X MEM, with 1X AA, 2mM L-glutamine, 2% FBS, 0.225% sodium bicarbonate and 0.75 % methylcellulose (Sigma M0152)) was added. Plates were incubated at 37°C with 5% CO_2_ for 6 days, checking daily for cytopathic effect. We then added 1 mL of staining solution (10% neutral-buffered formalin with 0.01% crystal violet) to each well and stained at RT in a drawer protected from light for 1-2 days. All wells were washed with sterile Milli-Q water twice before counting plaque forming units (PFUs).

### Bacterial growth conditions

*Dolosigranulum pigrum* strain KPL3065 (77) and *Streptococcus pneumoniae* strain 603 (78) were grown on BD BBL Columbia agar medium with 5% sheep’s blood (BAP) at 34°C in a humidified 5% CO_2_ incubator. To obtain single colonies, we struck *D. pigrum* from a frozen 15% glycerol stock stored at –80°C onto BAP. After 32 h of growth, 10-15 *D. pigrum* single colonies were collected with one sterile cotton swab (Puritan #25806) and bacteria were spread as a small lawn on a sterile 47-mm-diameter, 0.2-μm polycarbonate membrane (Millipore Sigma #GTTP04700) atop BAP, preparing 4 such plates. After 36-40 h of growth, we collected *D. pigrum* from 4 membranes using a sterile cotton swab, then resuspended the cells in EBSS. We struck *S. pneumoniae* from a frozen glycerol stock for singles directly onto BAP. After 9-10 h of growth, 10-15 colonies were picked up with a sterile cotton swab and swabbed as a lawn onto a sterile 47-mm, 0.2-micron polycarbonate membranes atop BAP, preparing 4 plates. After 15 h of growth, we gathered *S. pneumoniae* lawns from 4 membranes using a sterile cotton swab, resuspended cells in EBSS. We struck *H. influenzae* strain 86-028NP (79) from a frozen glycerol stock for singles directly onto a chocolate agar (Hardy Diagnostics #E14), preparing two plates. After 24 h of growth at 34°C in a humidified 5% CO_2_ incubator, 5-10 colonies were picked up with a sterile cotton swab and swabbed as a lawn onto a sterile 47-mm, 0.2-micron polycarbonate membranes atop chocolate agar, preparing 3 plates. After 24 h of growth, we collected the *H. influenzae* mini-lawn from one membrane, resuspended it in EBSS. The EBSS suspensions of each of the three bacteria were adjusted by optical density at 600 nm (OD_600_) to yield ∼1×10^7^ CFUs per 15 μL EBSS (as described in (39)).

### Bacterial colonization and viral infection of HNO-ALI

We colonized HNO-ALI with bacteria according to our methods in Boyd and Kafer et al. (39). Briefly, 15 μL of each bacterial suspension (∼1×10^7^ CFUs) was gently pipetted onto the surface of the corresponding HNO with 15 μL of EBSS alone pipetted onto uncolonized control HNO-ALI. The 24-well transwell plates were centrifuged at 120 x g for 1 min at RT in an Eppendorf Centrifuge 5430 R to help inoculate the bacteria into the apical mucus layer. HNO-ALI were incubated at 34°C in a humidified 5% CO_2_ incubator. After 23 h of bacterial colonization, basal medium was collected from each well and transferred to a microcentrifuge tube. After addition of fresh basal medium to each well, HNO-ALI were incubated at 34°C and 5% CO_2_ for 1 h. We then infected HNO-ALI with RSV/A/Ontario at an MOI of 0.01 (with a median of 5.9 x 10^3^ PFUs per HNO-ALI across all experiments) by diluting the viral stock in EBSS and adding 15 μL of the viral suspension to the apical side of the HNO. (This small volume was used to preserve the ALI and to avoid the need to remove excess inoculation liquid, which would wash away some of the bacteria.) RSV-uninfected control HNO-ALI wells received 15 μL of EBSS alone. We incubated HNO-ALI at 34°C and 5% CO_2_ for 2 days at which point spent basal medium was collected and fresh basal medium was added to all wells. HNO-ALI were incubated for 2 more days before collection at 4 dpvi (which was 5 days post bacterial colonization).

### Apoptosis assays

An apical wash and basal medium were collected from all HNO-ALI at 4 dpvi, including all controls, and immediately stored at –80°C. The apical wash consisted of two 200 μL rinses of the apical cell layer with AODM w/o HH, pipetting up and down three times with each rinse (400 μL total). To measure the amount of apoptosis in the apical wash, we used the Promega Caspase 3/7 Assay System (#G8091) according to manufacturer’s instructions. A standard curve was generated by making serial dilutions of recombinant Caspase-3 (Enzo Life Sciences BMLSE1695000) and measuring levels in the assay. This standard curve was used to calculate the level of caspase for each sample. The code for this calculation can be found at https://klemonlab.github.io/RSVBac_Manuscript.

### RNA extraction from HNO-ALI

Masterpure Complete DNA and RNA purification kit reagents (Biosearch Technologies #MGP04100) were used for RNA extraction. Four dpvi, basal medium was removed from each transwell then 600 µL of Tissue Lysis Solution with 2 µL of 50 µg/µL Proteinase K was added apically to each HNO followed several minutes later by pipetting up and down 10 times to remove the cell layer. Each cell suspension was transferred to 1.5 mL microcentrifuge tube and vortexed for 10 seconds (s). The cell suspension was then transferred to a Lysing Matrix B tube (MP Bio #6911100) for bead beating in a FastPrep-24 5G (MP Biomedicals Ver: 6005.3) for 45 s at 6.0 m/s 4 times, resting on ice for 30 s in between each step. Tubes were then incubated in a heat block at 65°C for 15 min, vortexing every 5 min. We then placed tubes on ice for 5 min before adding 350 µL of MPC Precipitation Reagent to each tube and vortexing for 10 s. Tubes were centrifuged at 15000 x g for 10 min at 4°C in an Eppendorf Centrifuge 5430R and supernatant was transferred to a clean 1.5 mL microcentrifuge tube (careful to avoid transfer of beads). The centrifuge step was repeated to remove all debris from the microfuge tube and supernatant was transferred to a clean 5 mL tube. To precipitate nucleic acids, 700 µL of isopropanol was added to each tube and mixed by inverting 40 times. We centrifuged tubes at 13,000 rpm for 10 min at 4°C before removing the isopropanol. The nucleic acid pellet was washed twice with 1000 µL of 70% ethanol, centrifuging at 13,000 rpm for 2 min at 4°C, before discarding the ethanol. The pellet was air-dried for approximately 1 h until it was translucent and no visible ethanol remained. The pellet was then dissolved in 100 µL nuclease-free water then treated with Turbo DNase (Invitrogen AM1907) by adding 10 µL of Turbo DNA-free buffer and 2 µL of DNA-free DNase to each tube, flicking to mix the reagents. Tubes were incubated at 37°C for 45 min, then another 2 µL of DNase was added and tubes were incubated for another 45 min. We then added 24 µL of inactivation reagent to each tube and incubated for 5 min at RT, flicking each tube to mix every 30 s. Tubes were centrifuged at 10,000 x g for 2 min at RT and the supernatant containing RNA was transferred to a clean tube then flash frozen in an ethanol-dry ice bath. RNA was stored at –80°C until sequencing.

### Transcriptomics

RNA samples were submitted to SeqCoast Genomics where libraries were prepared with the Illumina Stranded Total RNA Prep Ligation Kit with Ribo-Zero Plus Microbiome (#20072063) and indexed with Illumina Unique Dual Indexes. The Illumina NextSeq2000 platform was used for sequencing with the 300-cycle flow cell kit to generate paired reads (2 x 150bp). 1-2% PhiX spike-in was included to ensure accurate base calling. DRAGEN v4.2.7 was used for demultiplexing, adapter trimming, and run-level analytics. Individual sample quality was evaluated using FastQC, and summary metrics were inspected to confirm sequencing performance. RNAseq reads were aligned using STAR v2.7.10b (82) and gene expression was quantified using RSEM v1.3.3 (83). The Homo sapiens GRCh38 reference genome and corresponding annotation files were downloaded from Ensembl (soft-masked version, release 113) and used to generate index files for STAR (with ––sjdbOverhang 149) and RSEM. STAR was run with ––quantMode TranscriptomeSAM to generate transcriptome-aligned BAM files. After confirming that the percentage of aligned reads was consistent across samples, BAM files were randomly subsampled to match the sample with the lowest read count (95 million reads per sample) using samtools v1.17 (84). rsem-calculate-expression was run on the subsampled BAM files using ––estimate-rspd to improve quantification accuracy. Gene-level quantification results were imported into R using tximport (85), which produces length-scaled counts and an offset matrix for DESeq2. Transcripts with zero length were corrected to a length of one to avoid computational errors during normalization. Rarefaction curves were generated using the vegan package (86) to confirm that sequencing depth was sufficient to capture the majority of expressed genes. Gene-level counts and the associated length offset matrix from tximport were analyzed with DESeq2 (87). An initial model (∼ virus + species + line + seq_run) was used for normalization and quality assessment. Principal component analysis was performed on variance-stabilizing transformed (VST) data (87, 88) using the top 500 genes by variance to assess the primary sources of variation and evaluate potential batch effects. Differential gene expression analysis was performed using DESeq2 (87) with ashr log₂ fold change shrinkage (89). The model included virus status, bacterial species, their interaction, and HNO line: ∼ virus + species + virus:species + line. Individual contrasts were calculated (**Table S2-4**) for the effect of bacterial species-specific modulation of the epithelial response to viral infection (interaction terms). Gene set enrichment analysis (GSEA) (90) was performed using fgseaMultilevel() from the fgsea package (91). Genes were ranked by the Wald statistic from DESeq2, and enrichment was tested against Hallmark gene sets from the Molecular Signatures Database (MSigDB) (92) obtained via the msigdbr package (93) (**Table S5A**). Both, for differential expression and gene set enrichment, *p*-values were adjusted using the Benjamini–Hochberg false discovery rate (FDR) method (94) to account for multiple testing. For complete bioinformatic scripts, analysis code, R session information, and package versions see https://klemonlab.github.io/RSVBac_Manuscript.

### Immunoassays for detection of cytokines

Four days after RSV infection, basal medium was removed from each transwell and transferred to a 1.5 mL microcentrifuge tube then immediately frozen at –80°C. The apical side of each HNO was then washed twice with 200 μL of AODM and both washes were transferred to a single 1.5 mL microcentrifuge tube then immediately frozen at –80°C. Samples were submitted to the BCM Digestive Disease Center Functional Genomics and Microbiome core for immunoassay using the following Millipore Sigma magnetic-bead panel kits: 1) Human Cytokine Panel A (FGF-2/FGF-basic, MIG, IFNg, IFNa2, TNF-a, IP-10, RANTES, IL-1α, IL-1ra, IL-1β, IL-5, IL-6, IL-8, IL-10, IL-12(P70), IL-13, IL-18, MIP-1a, MIP-1b, MCP-1, MCP-3, IL-17A, IL-17E/IL-25, Eotaxin/CCL11, G-CSF, GM-CSF, VEGF-A); 2) Human Cytokine Panel II (IL-33, TRAIL, and TSLP; Human Cytokine Panel III with IL-29, I-TAC, CCL20/MIP-3a, and MIP-3b); 3) Human MMP Panel 2 (MMP1, MMP2, MMP7, MMP9, and MMP10); and 4) Human TIMP Panel (TIMP1). The core performed and analyzed immunoassays per the manufacturer’s instructions with Luminex xPONENT for Magpix (version 4.2, build 1324) on a Magpix instrument. Data were analyzed with Milliplex Analyst (version 5.1.0.0, standard build 10/27/2012). For downstream analysis, all values below the manufacturer’s limit of detection (LOD) were set to the LOD. The analyte FGF-2/FGF-basic was not able to be analyzed due to high background levels in the medium, and it was removed from all analyses, leaving a total of 39 cytokines. Also, due to low bead count in the multiplex ELISA, the following data points were removed from the analysis of the experiment on 05/01/2025 for HNO9007 4dpvi: 1) Hin+RSV apical IFNL1, CXCL11, CCL20, CCL19, and 2) Dpi+RSV apical IFNL1, CCL19. For each timepoint analyzed (0 and 4 days post-viral infection), if more than 25% of the samples for a given cytokine in a given compartment (apical vs. basal) were below the LOD, that cytokine-location combination was excluded from statistical analysis for that timepoint. Principal component analysis (PCA) and Partial Least Squares Discriminant Analysis (PLS-DA) were performed using the mixOmics package (95) to assess overall patterns of cytokine response. PCA was performed on mean-centered and scaled data with a multilevel design accounting for experimental date effects to remove batch variation. PLS-DA models were validated using 3-fold cross-validation repeated 50 times, with the optimal number of components selected based on the minimum balanced error rate (BER) using centroids distance classification (**Fig. S3**). See Statistical Analysis section for description of the univariate analysis with linear mixed-effects models (LMMs).

### Statistical Analysis

All analyses were performed in R (96) using RStudio (97). Data manipulation and visualization utilized the tidyverse suite (98) and related packages. Session information and version of all used packages is provided in https://klemonlab.github.io/RSVBac_Manuscript. For bacterial CFUs, RSV PFUs and copy number (RT-qPCR), caspase activity, VST-normalized expression for GSEA leading genes, and cytokine concentrations, we employed linear mixed-effects models (LMMs) using the lme4 package (99). All continuous response variables were log-transformed prior to analysis to satisfy model assumptions of normality and homoscedasticity. The general model structure included experimental condition (bacteria species and/or virus status, and their interaction when applicable) as fixed effects, with HNO line and experimental date as random effects to account for biological and technical variability. Data were stratified by HNO compartment location (apical vs. basal) and analyzed separately. In some analysis the model was extended to include time (dpvi or dpbc) as a fixed effect; in other cases, data was stratified an analyzed separately by time point. Detailed code and specific model designs for each analysis are available in https://klemonlab.github.io/RSVBac_Manuscript, and the results of the statistical analysis is summarized in Tables S1, S5B and S6. Model singularity was assessed for all fitted models; when singularity indicated negligible variance for nested random effects, models were refitted accordingly (usually by removing HNO line as a random effect). Statistical significance of fixed effects was evaluated using ANOVA with *p*-values computed using Satterthwaite’s approximation for degrees of freedom via the lmerTest package (100). Post-hoc pairwise comparisons (i.e., contrasts within each analysis) were conducted using estimated marginal means (EMMs) with the emmeans package (101). Within each individual analysis, *p*-values were adjusted using the Holm method to control family-wise error rate. Except for cytokine data, where *p*-values from all contrasts across all cytokine-location combinations were adjusted using the false discovery rate (FDR) method (94) to account for multiple testing across the large number of analytes. Model predictions were computed on the log-transformed scale, and approximate 95% confidence intervals were obtained using ±2 standard errors. Predictions and confidence intervals were then back-transformed to the original scale for visualization along data point representing individual biological replicates.

## Data Availability

All curated data (R-formatted.rds files) and analysis code (.qmd notebooks) required to reproduce the analyses and generate all figures are available at https://klemonlab.github.io/RSVBac_Manuscript. RNAseq sequencing data will be deposited in BioProject PRJNA1417248.

## Acknowledgements

We thank the donors who make HNO-ALI possible. We thank Lauren Bakaletz and Rick Malley for bacterial strains; Letisha Aideyan, Trevor McBride, Ashley Murray, and Gina Aloisio for guidance on viral protocols and assays; and members of the Respiratory Organoid Group and the Lemon Lab for helpful discussions. This research was supported in part by funding from the National Institute of Allergy and Infectious Diseases of the National Institutes for Health (NIH) under award U19AI157981 (K.P.L., S.E.B.), U19AI144297 (S.E.B., P.A.P.), U19AI116497 (P.A.P., S.E.B.) and by Baylor College of Medicine (via seed funds to K.P.L.) Services from the Texas Medical Center Digestive Disease Center Functional Genomics and Microbiome Core (FGM) and Tissue Analysis & Molecular Imaging Core (TAMI) were supported in part by NIH grant P30DK056338. Seed funding from Baylor College of Medicine (K.P.L.).

## Author Contributions

Conceptualization: L.A.K., K.P.L. Methodology: L.A.K., I.F.E., A.I.B., V.A., K.P.L., P.A.P. Investigation: L.A.K. Resources: A.R.T, A.K., S.E.B., I.F.E. Data curation: I.F.E., L.A.K. Formal analysis: I.F.E., L.A.K. Software: I.F.E. Validation: L.A.K., I.F.E. Visualization: L.A.K., I.F.E. Writing – original draft: L.A.K., I.F.E., K.P.L. Writing – review & editing: L.A.K., I.F.E., K.P.L., A.I.B., V.A., P.A.P., S.E.B., A.R.T, A.K. Funding acquisition: K.P.L., S.E.B., P.A.P. Supervision: I.F.E., K.P.L.

## Declaration of Interests

The authors declare no competing interests.

## Supplemental Information

Table S1. Microbial quantification and caspase statistics

Table S2. HNO-ALI DEGs: *S. pneumoniae*-RSV statistical interactions

Table S3. HNO-ALI DEGs: *H. influenzae*-RSV statistical interactions

Table S4. HNO-ALI DEGs: *D. pigrum*-RSV statistical interactions

Table S5. GSEA NES and selected VST statistical analysis

Table S6. Cytokine statistics

Fig S1. Bacteria-only CFUs and 1 dpbc cytokines

Fig S2. HNO-ALI DEGs: PCA and selected expression heatmaps

Fig S3. Cytokine PCA and PLS-DA performance

